# Stage-resolved Spatial Multi-omics Reveals Myeloid Niches in Human Atherosclerotic Plaques

**DOI:** 10.1101/2025.10.09.681522

**Authors:** Pengbo Hou, Huan Cai, Zhanhong Liu, Jiankai Fang, Ziyi Wang, Shiqing Wang, Qingkang Lyu, Peishan Li, Changshun Shao, Gerry Melino, Klaus Ley, Bo Hu, Yufang Shi

**Affiliations:** The Third Affiliated Hospital of Soochow University, Institutes for Translational Medicine, State Key Laboratory of Radiation Medicine and Protection, Suzhou Medical College of Soochow University, 215123 Suzhou, China; Vascular Surgery Department, The Fourth Affiliated Hospital of Soochow University, 215123 Suzhou, China; Department of Experimental Medicine, TOR, University of Rome Tor Vergata, 00133 Rome, Italy; Laboratory of inflammation Biology, Center for Autoimmune Disease, La Jolla Institute for Immunology, La Jolla, CA, USA; Department of Physiology, Immunology Center of Georgia, Augusta University, Augusta, GA, USA

## Abstract

**Background:** Atherosclerosis, a leading cause of heart attack and stroke, involves intricate immune cell dynamics within arterial plaques, yet their spatial organization and functional roles remain elusive.

**Methods:** We combined Visium HD spatial transcriptomics, imaging mass cytometry, and single-cell RNA-seq across fatty-streak, advanced, and restenotic plaques to map myeloid architectures and relate them to lesion geography and extracellular matrix features. Slingshot trajectory analysis resolved macrophages differentiation path. *In vitro*, we tested microenvironmental and lipid cues separately: fibronectin (FN) exposure during PMA-driven THP-1 differentiation and oxidized LDL (oxLDL)-induced foam cell formation in murine bone-marrow derived macrophages (BMDMs).

**Results:** Our analysis identified seven macrophage subsets and two neutrophil populations with distinct spatial distribution and functional roles. In early lesions, neutrophils expressing MMP9, MPO, p47phox, TGF-β1 and arachidonate 5-lipoxygenase (ALOX5), aligned with proteolysis, inflammatory processes, and endothelial-mesenchymal transition features. In advanced plaques, macrophage subsets exhibit specialized functions: Ki67^+^ proliferative macrophages localized near necrotic cores, sustaining local population; SPP1^+^ macrophages, enriched in lipid handling and tissue remodeling, are prone to apoptosis/ferroptosis, potentially promoting necrotic core expansion; and C3aR^+^ macrophages form antigen-presenting niches with elevated HLA-DR and CD74, engaging T cells possibly through CXCL12–CXCR4 signaling. Slingshot trajectories indicated progression from C3aR⁺ toward SPP1⁺ remodeling states concentrated at fibronectin-rich rims. *In vitro*, FN increased MMP9 and TIMP1 in THP-1-derived macrophages, consistent with FN imprinting remodeling features characteristic of SPP1⁺ macrophages *in situ*. Concurrently, oxLDL-treated BMDMs showed enhanced lipid-handling and remodeling modules consistent with the SPP1 program.

**Conclusions:** These findings define conserved myeloid niches and support a microenvironment-imprinting model that links ECM composition and lipid loading to macrophage state transitions, providing a framework for microenvironment-targeted therapies to stabilize plaques and mitigate cardiovascular risk.

## Introduction

Atherosclerosis, a lipid-driven chronic inflammatory disorder, is characterized by leukocyte infiltration and necrotic core development within large arterial walls^1^. Atherosclerotic cardiovascular diseases (ASCVD), including myocardial infarction and stroke, remain leading global causes of mortality^2–4^. Clinical trials like CANTOS^5^ (targeting IL-1β) and RESCUE^6^ (targeting IL-6) underscore the pivotal role of immune modulation in atherosclerosis, necessitating detailed spatial and phenotypic characterization of plaque immune cells to inform the development of therapeutic stragegies^7^.

Single-cell technologies, including scRNA-seq and cytometry by time of flight (CyTOF), have uncovered diverse immune and stromal populations in murine and human plaques^7–9^. Macrophages exhibit distinct phenotypes, including resident-like, foamy TREM2^hi^, and inflammatory subsets^10,11^, with recent studies identifying proliferative (Ki67^+^) and interferon-inducible macrophages, expanding known myeloid diversity^12^. Neutrophils, previously considered transient, are now recognized as key contributors to early atherogenesis via matrix-degrading proteases, oxidative enzymes, and proinflammatory lipid mediators^13,14^. However, such insights have largely been derived from dissociated tissues, which disrupt spatial context and limit our understanding of microanatomical organization and cell–cell interactions. Computational tools like CellPhoneDB and NicheNet, which infer cell communications based on ligand–receptor gene expression, are constrained by their lack of spatial resolution^15^. Recent spatial transcriptomic studies have advanced understanding of plaque cellular architecture. Bleckwehl et al. integrated multi-cohort scRNA-seq data with 10x Visium (55 μm spot-based) spatial transcriptomics to map coronary artery plaque atlas, revealing endothelial–immune cell interactions and smooth muscle cell phenotypic transitions^16^. Conversely, Pauli et al. (bioRxiv preprint, 2025) applied Xenium (0.2 μm resolution) targeted in situ transcriptomics in human carotid plaques, proposing an HMOX1^+^ inflammatory to TREM2^+^ lipid-handling macrophage differentiation trajectory. These studies, while insightful, were limited by low resolution, restricted gene panels, and lacked protein-level validation. Imaging mass cytometry (IMC), with 1 μm^2^ resolution using metal-tagged antibodies, enables multiplexed, quantitative protein imaging, eliminating spectral overlap and tissue autofluorescence while preserving architectural context^17^. Widely applied in oncology and immunology, IMC supports biomarker discovery and niche-level cellular decoding^7,18^. Visium HD spatial transcriptomics (2 μm × 2 μm bin size) provides near single-cell resolution, enabling comprehensive, unbiased spatial gene expression profiling within intact tossue^19^.

In this study, we integrated Visium HD spatial transcriptomics, offering near single-cell resolution, with IMC for high-resolution protein imaging and scRNA-seq to delineate functionally specialized immune cell niches driving plaque initiation, progression, and restenosis. We identified distinct macrophage and neutrophil subsets, mapped their spatial distribution, and characterized niche-specific intercellular signaling networks shaping plaque evolution. While macrophage heterogeneity is well established, whether their phenotypes are lineage-intrinsic or microenvironmentally reprogrammed remains unclear. Our integrative spatial framework addresses this question and demonstrates IMC’s utility in decoding immune remodeling in human atherosclerosis.

## Methods

### Data Availability

A full description of materials and methods is provided in the Supplemental Material; see also the Major Resources Table in the Supplement. All animal and human studies were approved by the local ethics committees of the Experimental Animal Center, Soochow University, and the Fourth Affiliated Hospital of Soochow University. All human participants provided written informed consent under protocols approved by the respective institutional review boards.

Public single-cell RNA-seq datasets were obtained from the Plaque Atlas (Plaque Atlas — CZ CELLxGENE Discover) and Gene Expression Omnibus (GSE159677). The raw Visium HD spatial transcriptomics, bulk RNA-seq, and imaging mass cytometry datasets generated in this study will be deposited in a public repository and made available upon publication. Processed data and analysis code are available from the corresponding author upon reasonable request.

## Results

### Visium HD spatial transcriptomics identifies specialized macrophage niches in advanced plaques

To construct a detailed spatial immune atlas of human atherosclerosis, we analyzed 28 FFPE plaque specimens from carotid (n = 13) and lower extremity (n = 15) arteries (Supplementary Table 1), encompassing fatty streaks (n = 4), advanced plaques (n = 19), and restenosis lesions (n = 5). Von Kossa and H&E staining verified varying degrees of calcification at different stages (Figure 1A-B). We performed imaging mass cytometry (IMC) on serial sections to characterize single-cell phenotypes at high spatial resolution. We also performed Visium HD spatial transcriptomics on one typical advanced calcified carotid plaque to explore its gene expression patterns with spatial detail (Figure 1A).

**Figure 1.**
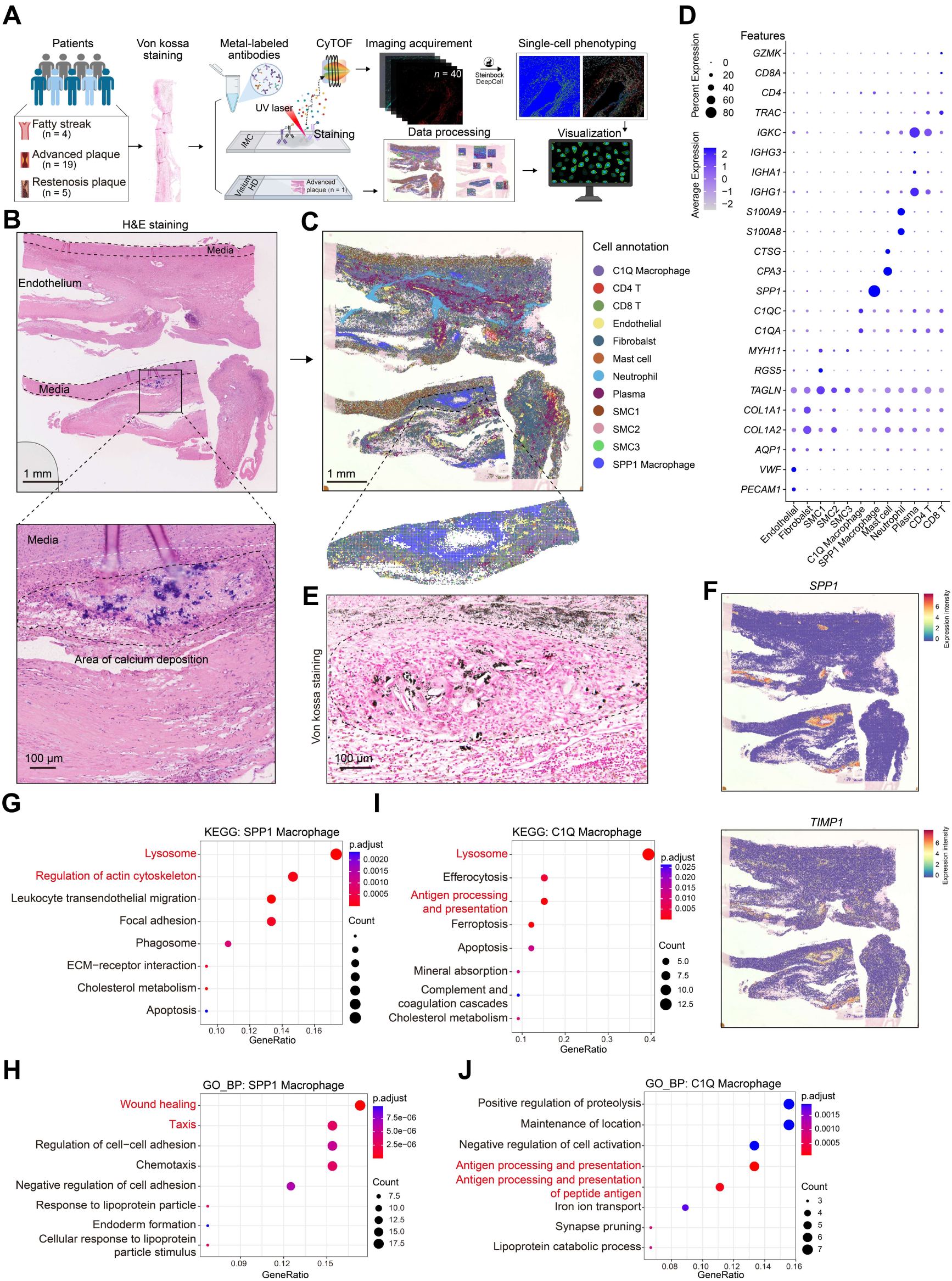
Visium HD spatial transcriptomics identifies specialized macrophage niches in human atherosclerotic plaques. **A,** Schematic workflow of plaque sample processing and analysis. FFPE plaque sections from fatty streak (n = 4), advanced plaque (n = 19), and restenotic plaque (n = 5) stages underwent Von Kossa staining, metal-labeled antibody staining for IMC, imaging acquisition, and single-cell phenotyping. **B,** Representative H&E staining of an advanced carotid plaque showing calcified regions within the intima and media. Scale bar, 1 mm (top image) and 100 µm. **C,** Spatial transcriptomic clustering identifies 12 distinct cell populations within advanced plaque tissue. Scale bar, 1 mm (top image). **D,** Dot plot showing expression of representative marker genes across the identified cell populations. **E,** Von Kossa staining confirming calcium deposition areas, co-localizing with regions enriched in SPP1 macrophages. Scale bar, 100 µm. **F,** Spatial feature plots showing expression of *SPP1* and *TIMP1* within the plaque section. **G–H,** KEGG (G) and GO Biological Process (h) pathway enrichment analyses for SPP1 macrophage subset genes. **I–J,** KEGG (I) and GO Biological Process (J) enrichment analyses for C1Q macrophage subset genes.

Histology revealed significant calcium buildup in the intima of advanced plaques (Figure 1B). Spatial transcriptomics pinpointed 12 unique cell groups using standard marker genes (Figure S1A), including two macrophage types: SPP1 macrophages (also known as foamy/TREM2^hi^ macrophages)^20^ and C1Q macrophages (co-expressing *C1QA*, *C1QC*, and *C3AR1*)—plus three smooth muscle cell variants (SMC1–3), endothelial cells (*PECAM1*), fibroblasts (*COL1A1*), neutrophils (*S100A8/9*), CD4^+^ and CD8^+^ T cells, plasma cells (*IGKC*), and mast cells (Figure 1C-D, Figure S1B). Spatial patterns of cell-type markers are shown in Figure S1C. H&E-based annotation outlined viable non-necrotic areas, necrotic cores, T cell-rich zones, ECM-rich areas, and regions with neovascularization/microvasculature, helping frame compartment-specific cell distributions and interactions (Figure S1C-D).Public scRNA-seq datasets (Plaque atlas from Traeuble et al^9^. and GSE159677) validated these cell groups, with SPP1 macrophages expressing *CD68*, *SPP1*, *MMP9*, *TIMP1*, and *Vimentin*, and C1Q macrophages featuring *C1QA*, *C1QC*, *HLA-DRA*, and *C3AR1* (Figure S2A-C and S3A-C). This cross-validation underscores the reliability of our Visium HD spatial transcriptomic data. Spatially, SPP1 macrophages mainly occupied calcified zones, implying involvement in lesion calcifcation^10^, while C1Q macrophages bordered these areas, creating an organized spatial niche (Figure. 1C). Von Kossa staining verified SPP1 macrophage alignment with calcification spots (Figure. 1E). Functionally, SPP1 macrophages strongly expressed *SPP1* (Osteopontin, OPN) and *TIMP1*^21^ (tissue inhibitor of metalloproteinases 1) (Figure. 1F), indicating roles in extracellular matrix (ECM) restructuring. KEGG pathway enrichment analysis showed SPP1 macrophage genes enriched in wound healing, cell migration, lysosomal function, and actin cytoskeleton regulation pathways (Figure 1G-H, Figure S2D and S3D). Conversely, C1Q macrophage genes were linked to antigen presentation and efferocytosis, aligning with scRNA-seq findings (Figure. 1I-J, Figure S2D and S3D). These results highlight unique spatial segregation and functional roles of macrophage subtypes in advanced human atherosclerotic plaques.

### IMC analysis uncovers macrophage and neutrophil diversity across plaque phases

To comprehensively map immune and stromal cells in human atherosclerotic plaques, we created a 40-marker IMC panel focusing on lineage and functional proteins (Figure 2A, Supplementary Table 2). Typical spatial layouts of key lineage and functional markers are displayed in Figure 2B and Figure S4A-B. With this panel, we assessed 267,467 single cells from 152 regions of interest (ROIs), detecting 30 unique cell clusters, such as macrophages, neutrophils, SMCs, endothelial cells (EC), fibroblasts, T cells, and mast cells (Figure 2C-D). Unsupervised clustering identified seven macrophage subtypes with distinct traits. SPP1⁺ macrophages, with elevated Vimentin, CD44, FAP, and CD36, displayed a foam cell-like profile with ECM remodeling abilities. C3aR⁺ macrophages had increased HLA-DR, suggesting antigen-presentation roles; Ki67⁺ macrophages showed proliferation indicators, implying local renewal to maintain macrophage numbers^22^. These proliferating “macrophages” could be newly infiltrated monocytes or early macrophages. We also found two CD38⁺ macrophage groups: CD38⁺ Mac and CD38⁺SPP1⁺ Mac. CD38, an NAD-depleting enzyme, promotes pro-inflammatory and senescent macrophage states^23–25^. Both groups had high phosphorylated p65 (pp65), signaling NF-κB activation. The CD38⁺SPP1⁺ macrophages notably co-expressed high SPP1 and the antioxidant factor Nrf2, indicating engagement in oxidative stress responses alongside inflammation. Other groups included TGF-β1⁺ macrophages (FAP^+^), possibly aiding fibrotic changes, and fibronectin-positive (FN^+^) macrophages (FN⁺ Mac), a small group likely involved in ECM modulation (Figure 2E).

**Figure 2.**
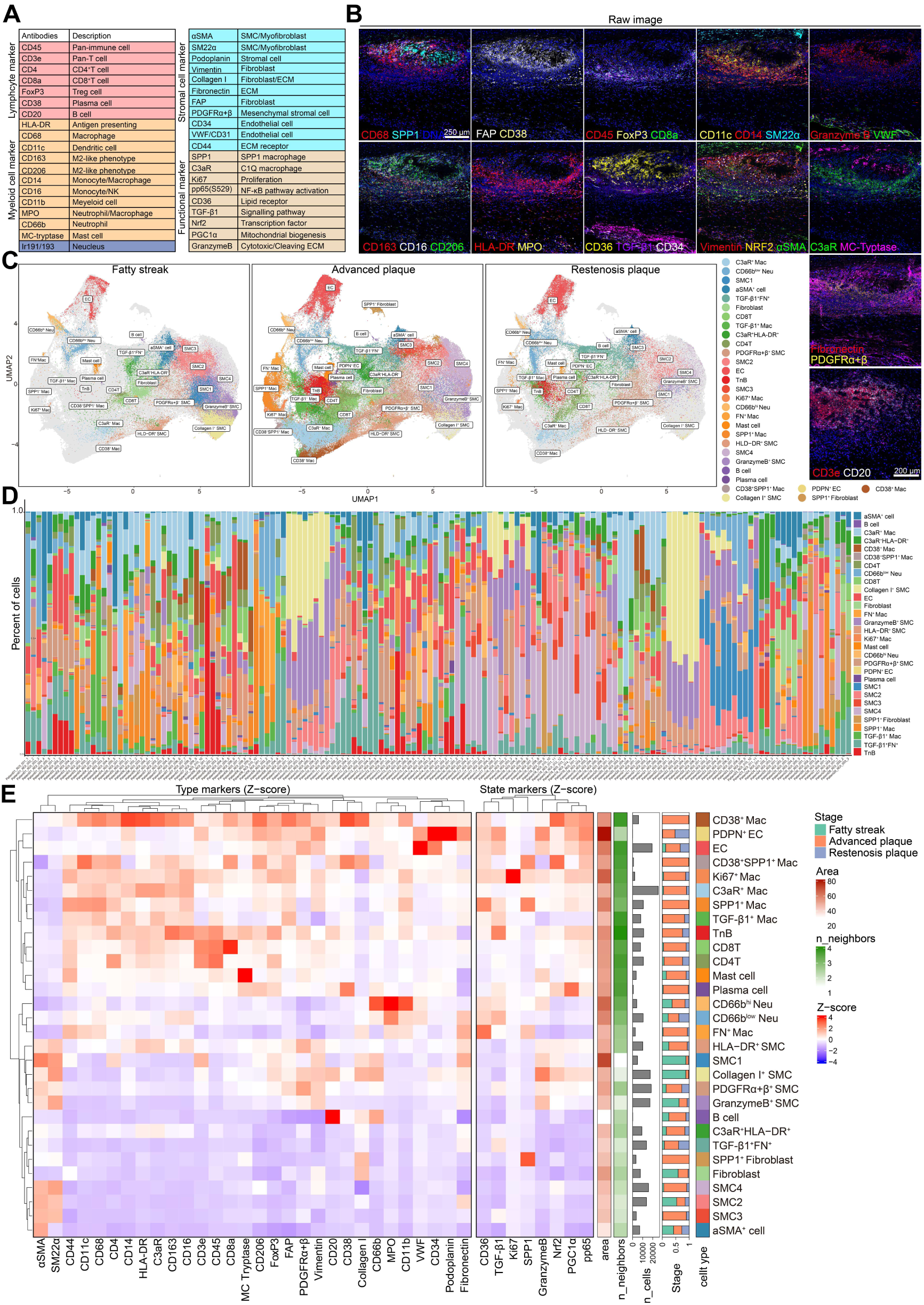
Imaging mass cytometry reveals macrophage and neutrophil heterogeneity across human plaque stages. **A,** Overview of the 40-metal conjugated antibody panel used for IMC profiling, targeting markers of stromal cells, myeloid cells, lymphocytes, and functional states. **B,** Representative multiplexed IMC images showing marker expression and spatial localization of key immune and stromal populations. Scale bars, 250 µm. **c,** UMAP clustering of 267,467 single cells from 152 ROIs across various plaque progression stages, identifying 30 distinct clusters annotated based on canonical marker expression. **D,** Bar plot showing the proportion of each cell type cluster in each ROI from different patients. **E,** Heatmap of type and state marker expression across identified cell clusters, with annotations showing cluster identity, stage distribution (fatty streak, advanced plaque, restenosis), cell counts and neighborhood area.

Two neutrophil subtypes emerged: CD66b^low^ neutrophils, expressing TGF-β1 and fibronectin but no granzyme B, and CD66b^high^ neutrophils, expressing granzyme B, MPO, CD36, and TGF-β1, suggesting stronger cytotoxic or NETosis potential. Both expressed CD36, a high affinity receptor for long chain fatty acids. While macrophage CD36 drives oxLDL uptake and foam cell formation^26,27^, earlier work showed neutrophils boost LDL buildup and oxidation via MPO-derived oxidants^13,14^. Our data build on this, revealing high CD36 expression in neutrophils and suggesting their contribution to lipid oxidation, foam cell development, and inflammation escalation. To evaluate shifts in immune and stromal makeup by disease phase, we grouped IMC-profiled cells by plaque type and performed UMAP embedding for fatty streak, advanced, and restenotic lesions (Figure 2C and 2F). Fatty streaks featured heavy neutrophil infiltration—especially CD66b^hi^ and CD66b^low^subsets—with low macrophage and lymphocyte levels, emphasizing early neutrophil roles. In contrast, advanced plaques displayed substantial expansion of macrophages, including SPP1⁺, C3aR⁺, CD38⁺, CD38^+^SPP^+^, FN^+^, and Ki67⁺ subtypes, plus more fibroblasts and lymphocytes, indicating immune activation and matrix changes. Restenotic plaques had less myeloid variety but more fibroblasts and PDPN⁺ endothelial cells, hinting at post-interventional fibrotic shifts.

To probe anatomical heterogeneity, we divided advanced plaques by vascular site. Aligning with reports that femoral plaques are less inflammatory than carotid plaques^28^. Carotid lesions had more foamy, inflammatory, and proliferative macrophages like C3aR⁺, CD38⁺, Ki67⁺, and SPP1⁺. Femoral plaques showed higher B cells and TGF-β1⁺ macrophages (Figure S5A-C). These findings show site-specific immune settings, with carotid plaques favoring inflammatory niches and femoral plaques, leaning toward fibrotic or regulatory profiles.

### HOCl-induced fibronectin oxidation colocalizes with neutrophil niches and TGF-β1 during early atherogenesis

To define neutrophil spatial behaviors and functional states across plaque stages, we combined IMC and transcriptomic data. In fatty streaks, IMC showed mainly CD66b^low^ neutrophils near the subendothelial lumen, with rare CD66b^hi^ neutrophils on the endothelium (Figure 3A). Interestingly, the endothelial marker vWF was attenuated in these regions, accompanied by prominent TGF-β1 and fibronectin (FN) signals, consistent with early EndMT-like features^29,30^. Feature plots of MPO and FN (Figure 3B) highlighted FN-dense settings around neutrophil-rich areas. Because tyrosine residues are susceptible to oxidation by hypochlorous acid (HOCl) generated by neutrophil MPO^14,31^, immunohistochemistry was used to examine oxidative marks: HMOX1-high areas overlapped with 3-chlorotyrosine (Figure 3C), a sign of HOCl-associated protein modification in plaques^32^. These zones also showed MPO⁺ neutrophils with FN accumulation (Figure 3D), possibly heightening oxidative niches coincide with matrix-remodeling hotspots and may facilitate early EndMT. Single-cell RNA-seq indicated that plaque neutrophils up-regulate *HMOX1* and express *TGFB1* (Figure 3E). IMC further showed spatial overlap of TGF-β1 with neutrophil markers and minimal overlap with smooth-muscle (SM22α/αSMA) or endothelial (vWF) markers (Figure 3F-G), suggesting that neutrophils could contribute to local TGF-β1 in early lesions. While alternative sources cannot be excluded, the observed vWF loss together with FN deposition is consistent with a model in which neutrophil-associated TGF-β1 may promote EndMT at fatty-streak.

**Figure 3.**
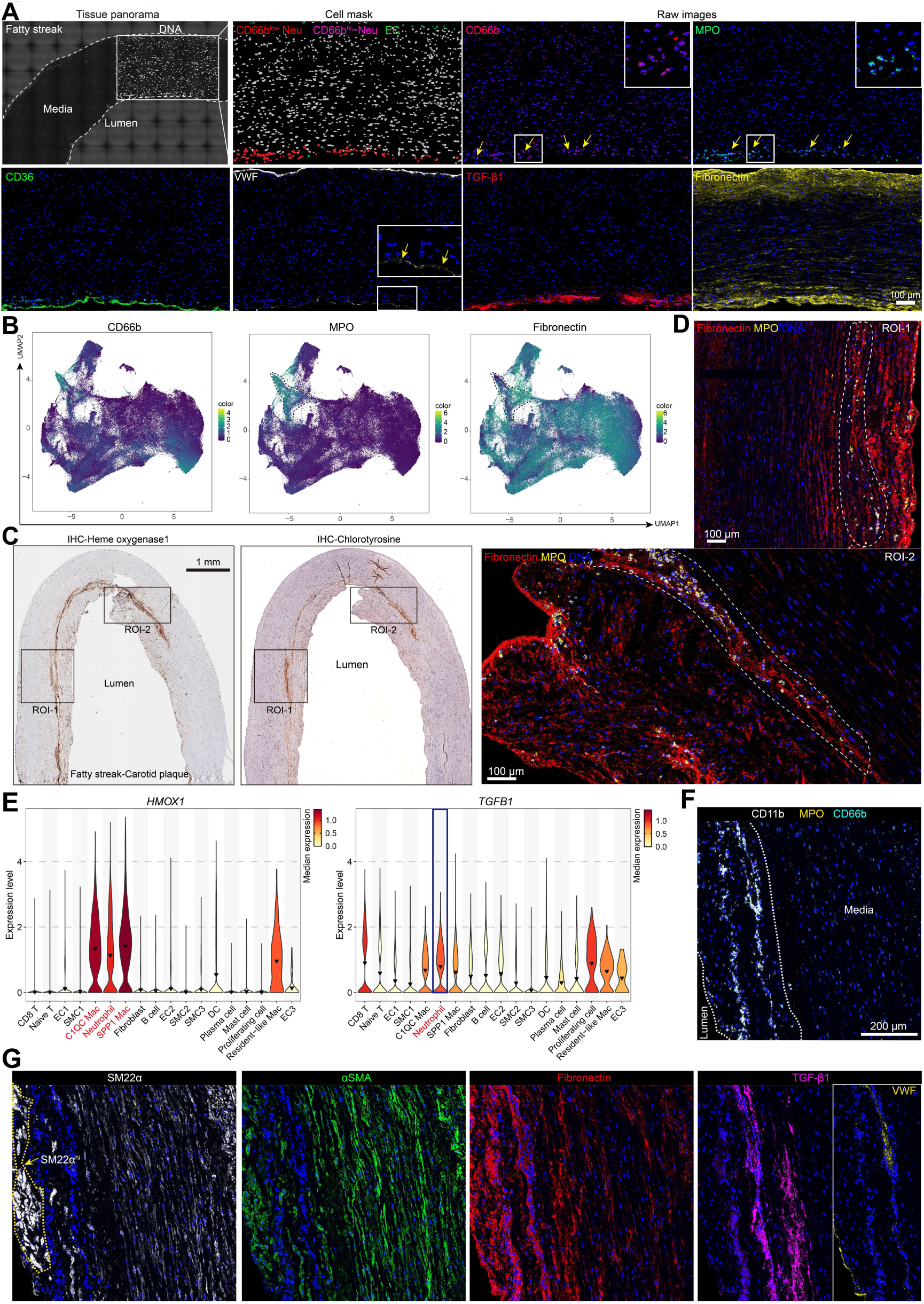
Neutrophil bands at the luminal rim associate with TGF-β1, EndMT and FN modification. **A,** IMC of a fatty-streak carotid plaque (tissue panorama and cell mask). Neutrophils (CD66b⁺/MPO⁺) concentrate along the luminal intima and inner media (yellow arrows), juxtaposed to FN-dense lamellae and TGF-β1 signal; endothelial cells (vWF⁺) line the lumen and CD36 marks lipid-handling cells. Scale bar, 100 µm. **B,** IMC-derived single-cell UMAPs showing the distributions of CD66b, MPO, and FN intensities across the section; neutrophil clusters co-localize with FN-rich territories. **C,** IHC validation on serial sections demonstrates oxidative stress in fatty-streak lesions: HMOX1 (heme oxygenase-1) and 3-chlorotyrosine. Scale bar, 1 mm. **D,** Higher-magnification IMC confirms spatial apposition of FN (red) and MPO⁺ neutrophils (Yellow) at representative ROIs near the lumen and shoulder. Scale bar, 100 µm. **E,** Violin plots of HMOX1 and TGFB1 across annotated cell types reveal elevated oxidative-stress and TGF-β programs in neutrophils and selected stromal/immune populations. **F,** IMC for CD11b, MPO, and CD66b localizes neutrophil bands to the intima–media interface (dashed lines). Scale bar, 200 µm. **G,** Structural context: SM22α and αSMA delineate smooth-muscle layers; FN forms parallel sheets beneath the endothelium; TGF-β1 tracks with neutrophil bands rather than endothelial cells, consistent with EndMT-permissive conditions and potential neutrophil-driven FN modification along the luminal surface.

### Neutrophils display phase-specific spatial patterns and oxidative activities

In advanced plaques, CD66b^low^ neutrophils spread through the intima and gathered near necrotic cores, with little presence in intact endothelial areas (Figure 4A). Neutrophils appeared in tertiary lymphoid structures (TLS), often near adventitial small vessels (Figure 4B), suggesting involvement in antigen display or inflammation control. In restenotic plaques, many CD66b^low^ neutrophils clustered at endothelial damage sites, marked by low vWF, while CD66b^hi^ neutrophils stayed mostly on endothelium (Figure 4C-D). These areas had rich fibronectin and granzyme B. Since neutrophil granzyme B cleaves ECM like fibronectin and vWF expression, our results suggest neutrophils aid post-injury matrix restructuring and endothelial disruption in restenosis. Across multiple restenotic specimens with abundant neovascularization, neutrophils consistently localized adjacent to vWF⁺/CD34⁺ microvessels, with perivascular CD66b^low^ neutrophils; this spatial coupling— together with local TGF-β1, pp65 and fibronectin enrichment—is compatible with neutrophil involvement in endothelial plasticity (EndMT-like changes) and perivascular inflammation (Figure S6A-B). Overall, CD66b downregulation after migration signals activation, implicating neutrophils’ role in early oxidation, ECM breakdown, and localized inflammation across plaque types.

**Figure 4.**
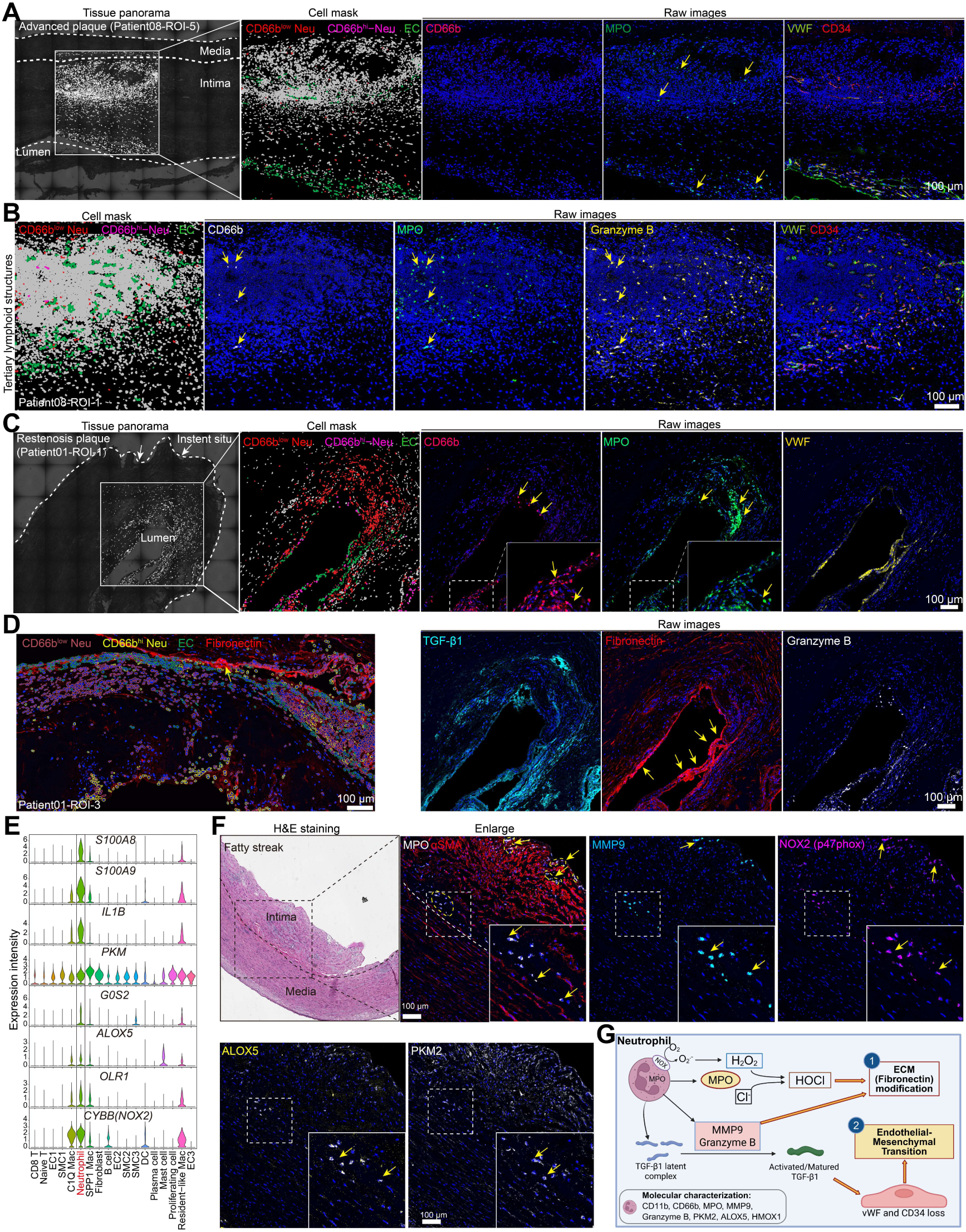
Stage-specific neutrophil localization and pro-oxidative programs across human atherosclerotic plaques. **A–C**, IMC-based in situ profiling of neutrophils in an advanced plaque (A), a region containing a tertiary lymphoid structure (B), and a restenotic plaque (C). Left: tissue panoramas with ROIs; middle: cell masks showing CD66b^hi^ (magenta) and CD66b^low^ (white) neutrophils, endothelial cells (vWF⁺, green); right: raw channels (CD66b, MPO, vWF/CD34, FN, granzyme B) illustrating spatial localization relative to lumen, intima, and media. Scale bars, 100 µm. **D**, Composite IMC places CD66b/MPO neutrophils within FN-rich rims and areas with TGF-β1 and granzyme B signal in restenotic tissue. Scale bar, 100 µm. **E,** Violin plots (cell-type annotations from calcified human carotid plaques, GSE159677) show neutrophil-enriched genes (*S100A8*/*A9*, *IL1B*, *PKM*, *G0S2*, *ALOX5*, *OLR1*, *CYBB*/*NOX2*), consistent with oxidative/inflammatory and lipid-handling programs. **F,** H&E staining image and IMC validation (right) in fatty-streak lesions demonstrate high MPO, MMP9, NOX2 (p47phox), ALOX5, and PKM2 within subendothelial, transmigrated neutrophils. Scale bars, 100 or 200 µm. **G,** Mechanistic schematic: neutrophils generate ROS via NOX2 and convert H₂O₂ to hypochlorous acid (HOCl) via MPO, which modifies FN and other ECM proteins; together with MMP9 and granzyme B, these processes promote activation of latent TGF-β1, creating EndMT-permissive conditions at the luminal rim in early plaques.

Though IMC detected neutrophils well, spatial transcriptomic profiling via Visium HD transcriptomics found a S100A8/9^+^ neutrophil group, but classic markers like CD66b (CEACAM8) and MPO had low RNA expression— possibly from FFPE RNA loss or poor transcript-protein match. The S100A8/9^+^ cluster did not align spatially with CD66b^+^ or MPO^+^ neutrophils from IMC analysis, showing method discordance (Figure S6C). Also, some scRNA-seq studies label S100A8/9^+^ cells as IL1B^+^ macrophage subsets^33^, blurring lineage. Analyzing public scRNA-seq (GSE159677) from carotid plaques, we found a unique S100A8/9^+^ neutrophil group enriched in phagosome and ROS pathways (Figure S6D). These cells co-expressed *IL1B*, *PKM2*, *G0S2*, *ALOX5*, *OLR1*, and *CYBB* (encoding the catalytic core of the NADPH oxidase 2 (NOX2) complex) (Figure 4E), marking activated, glycolytic, and oxidative state. IMC confirmed this proteomically, with migrated neutrophils high in MPO, MMP9, p47phox (NOX2 subunit for respiratory burst), ALOX5, and PKM2 (Figure 4F), backing roles in matrix breakdown, ROS generation, eicosanoid production, and metabolic shifts. These integrated results show spatial proteomics’ in neutrophil state definition, revealing distinct subsets with context-specific functions in oxidative damage, ECM changes, and immune regulation during atherosclerosis progression (Figure 4G).

### Macrophage spatial layout mirrors transcriptomic niches

To detail the architecture of macrophages within advanced plaques, we used IMC on serial sections from the same patient analyzed by Visium HD. Von Kossa showed calcium mainly in media, fibrous cap, and necrotic core (Patient 8-ROI5) (Figure 5A). Remarkably, SPP1^+^ and Ki67^+^ macrophages were near necrotic cores, while C3aR^+^ macrophages (C1Q type) created a surrounding “buffer” around SPP1^+^ ones, matching the Visium HD patterns (Figure 5B). This setup was consistent across advanced plaques from various patients (Figure 5C), indicating conserved macrophage niches. By contrast, CD38^+^ and TGF-β1^+^ macrophages lacked fixed localizations, suggesting greater microenvironment variability (Figure. S7A).

**Figure 5.**
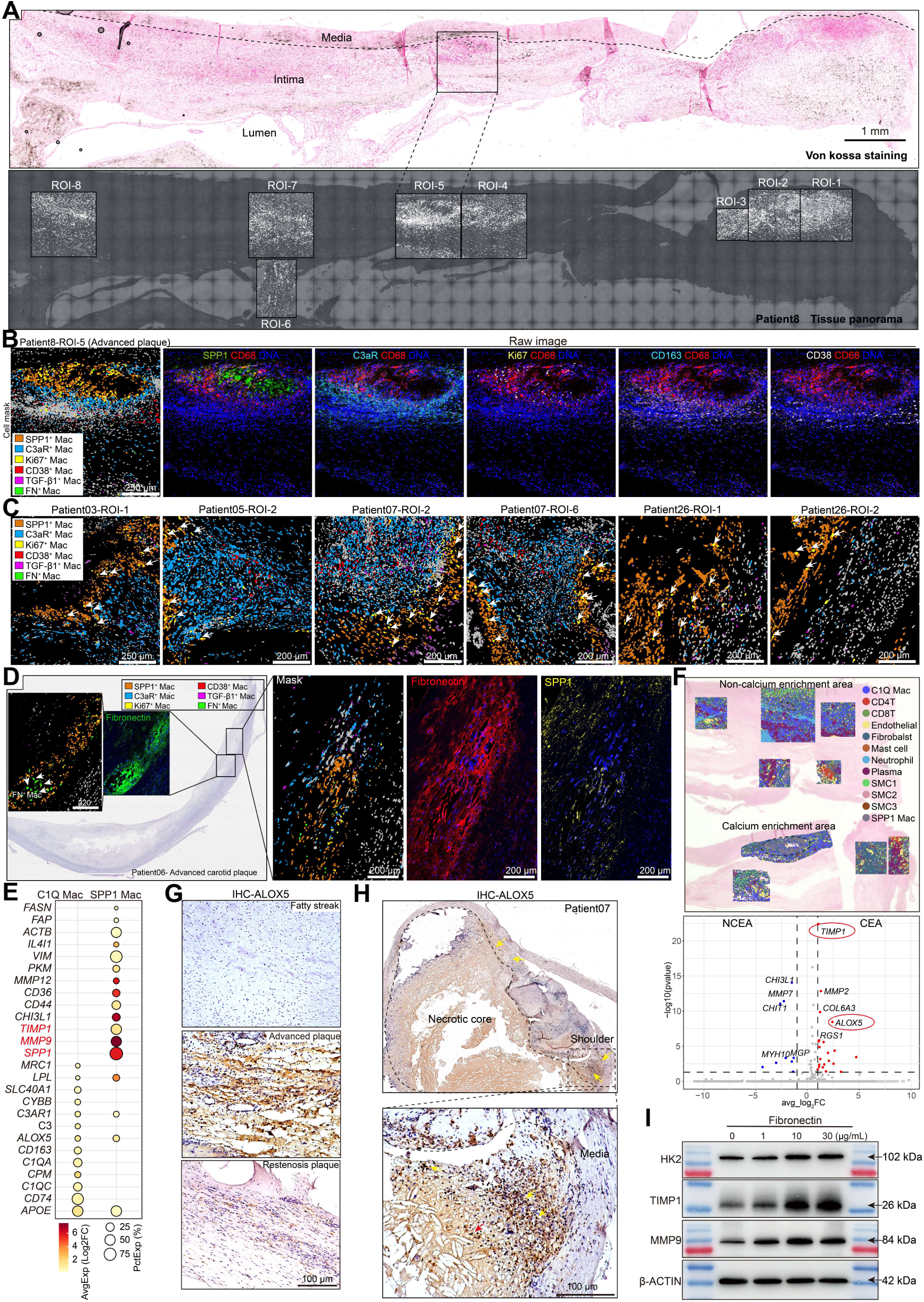
Distinct spatial organization and molecular signatures of macrophage subsets in advanced human plaques. **A,** Von Kossa staining of an advanced carotid plaque showing extensive calcium deposition within the intima and media. Scale bar: 1mm. Tissue panorama indicating ROIs selected for IMC analysis across different regions of the plaque. **B,** Cell mask of patient08-ROI-5 (advanced plaque), showing spatial distribution of macrophage subsets: SPP1^+^ (orange), C3aR^+^ (cyan), FN^+^ (green), CD38^+^ (red), Ki67^+^ (yellow), and TGF-β1^+^ macrophages (purple). Scale bar, 250 µm. **C,** Spatial distribution of macrophage subsets across different ROIs within the plaque. Scale bar, 200 or 250 µm. **D,** Whole-section view of a representative advanced carotid plaque with regional zoom-ins highlighting Fibronectin-rich areas populated by FN^+^ and SPP1^+^ macrophages. **E,** Dot plot comparing gene expression profiles of SPP1 and C1Q macrophages, showing enrichment of matrix remodeling genes (e.g., *TIMP1*, *MMP9*, *VIM*, *FAP*) in SPP1 macrophages and antigen presentation genes (e.g., *CD74*, *C1QA*, *C1QC*) in C3AR1 macrophages (Visium HD-annotated cell cluster). **F,** Differential gene expression analysis comparing SPP1 macrophages in calcium enrichment areas (CEA) versus non-calcium enrichment areas (NCEA). **G,** Immunohistochemistry showing ALOX5 expression in fatty streak, advanced, and restenotic plaques. Scale bar, 100 µm. **H,** ALOX5 expression in an advanced unstable plaque. Scale bar, 100 µm. **I,** Western blots detecting MMP9, TIMP1, and HK2 in PMA-induced THP-1 cells differentiating into macrophages in the presence of graded FN (0, 1, 10, 30 μg/mL). β-ACTIN serves as the loading control; approximate molecular weights are indicated at right.

Notably, FN^+^ macrophages clustered near necrotic cores, often with SPP1 expression (Figure 5D), in fibronectin-dense areas (Figure 5D, Figure S7B-C). Visium HD and public scRNA-seq confirmed high FN1 expression in SPP1^+^ macrophages (Figure S1C, S3B–C), showing SPP1⁺ macrophages add to ECM beyond typical sources like SMCs, fibroblasts, and ECs in late plaques.

Visium HD gene comparison showed distinct transcriptional patterns: C1Q macrophages (C3aR⁺ in IMC) had high *CD74*, *CPM*, *CD163*, and *C3*, for antigen presentation and complement involvement. SPP1⁺ macrophages upregulated genes associated with matrix remodeling (*MMP9*, *MMP12*, *TIMP1*, *CHI3L1*), lipid metabolism (*CD36*, *FASN*, *ALOX5*), and cytoskeletal dynamics (*VIM*, *FAP*, *CD44*) genes, plus *IL4I1* (Figure 5E). Markers like CD36, FAP, VIM, and CD44 were IMC-confirmed (Fig. 2f). Spatial analysis showed SPP1⁺ subtype variation: some in non-calcified areas (Figure 5F, top), others in calcified necrotic cores with higher *TIMP1*, *MMP2*, and *ALOX5* expression (Figure 5F, bottom). Immunohistochemistry verified strong ALOX5 in advanced plaques, with nuclear localization hinting arachidonic acid pathway activation. As leukotrienes from this pathway boost inflammation, ALOX5 likely aids local immune activation^34,35^. ALOX5 was enriched in unstable plaque shoulders but low in fatty streaks and restenotic lesions (Figure 5G-H).

Given the spatial co-enrichment of FN and SPP1⁺ macrophages, and the remodeling/glycolytic signature of the latter, we hypothesized that FN helps shape the SPP1^+^ macrophage phenotype. To test this, we added graded doses of FN during PMA-driven differentiation of THP-1 monocytes. FN induced a dose-responsive increase in MMP9 and TIMP1, and enhanced the glycolytic enzyme HK2, with a clear effect at 10 µg/mL (Fig. 5I). By contrast, applying FN to fully differentiated THP-1 macrophages did not significantly alter MMP9, TIMP1, or HK2 (Figure S7D), indicating that FN’s effect is stage-dependent and most pronounced during monocyte-to-macrophage differentiation. These data support a model in which FN in the plaque microenvironment biases monocyte–macrophage differentiation toward a controlled remodeling program (MMP9 and TIMP1) coupled to glycolytic re-wiring (HK2).

Collectively, macrophage subtype with matrix remodeling, glycolytic, and lipid-handling programs, cluster at necrotic cores rims, likely adding fibrotic and pro-inflammatory elements. C3aR⁺ macrophages with high expression of complement and antigen presentation machinery, may coordinate adaptive immunity at plaque edges. The interplay between ECM composition, particularly FN and macrophage state provides a mechanistic link between tissue remodeling and immune activation during plaque progression.

### SPP1^+^ and C3aR^+^ macrophages exhibit distinct metabolic, death, and epigenetic traits related to foam cell reprogramming

Since macrophage phenotypes and death modes influence plaque inflammation and progression, we designed IMC panel 2 (Figure 6A, Supplementary Table 2) based on transcriptional cues, assessing phagocytosis, lipid peroxidation, and death pathways. Markers included foam cell metabolism (PKM2, CD36, CPT1A, ACSL4, ALOX5), efferocytosis (MerTK), apoptosis (Caspase1, Cleaved caspase-3), necroptosis (pRIPK1), ROS (p47phox), and epigenetics modification (H3K9me3) (Figure 6A). From panel 1 heatmaps, major macrophage subsets- SPP1^+^, C3aR^+^, CD38^+^, CD163^+^—were readily recognized (Figure 6B-C. Over 124 ROIs from 15 patients, we profiled 249,027 cells (Figure S8A-B). SPP1^+^ macrophages had high CXCL12, MMP9, Caspase1, cleaved caspase3, PKM2, ALOX5, H3K9me3, CPT1A, p47phox; C3aR^+^ macrophages had lower levels except comparable ACSL4, ALOX5, p47phox, H3K9me3 (Figure 6B and 6D, Figure S9A). MerTK was low in SPP1⁺ and C3aR⁺, hinting poor efferocytosis, unlike CD163⁺ (in mixed TnB cluster) and MPO⁺ macrophages with higher MerTK for cell clearance (Figure 6C). Spatial views placed SPP1^+^ around necrotic cores, even without calcification (Figure 6D). Their poor efferocytosis and death profile likely fuel secondary necrosis. Necrotic centers had high pRIPK1, MPO, Caspase1, PKM2 (Figure 6D), showing myeloid death and metabolism. Immunohistochemistry showed RIPK3 at core edges and cleaved PARP1 around cores (Figure S10A-B), confirming necroptosis and apoptosis. In the same regions, high FASN (Fatty acid synthase) expression was also detected, reinforcing the lipid metabolic bias in SPP1^+^ macrophage co-localizing with death-prone niches (Figure S10A-C). H3K9me3, critical for transcriptional repression and chromatin compaction^36,37^, was high in SPP1^+^, C3aR^+^, CD163^+^, but absent in CD38^+^ macrophages, suggesting epigenetic control of macrophage functional states^38^ (Figure 6D). Given the spatial proximity, C3aR^+^ (C1Q) macrophages may shift to SPP1^+^ macrophages in necrotic niches. Based on public scRNA-seq data, Slingshot trajectory analysis revealed a differentiation trajectory from C1Q macrophages to SPP1 macrophages (Figure 6E), accompanied by downregulation of C1Q family members and antigen-presentation genes (*C3AR1*, *HLA-DRA*, *HLA-DRB1*, *CD74*) and upregulation of *VIM*, *FN1*, *PKM*, and *SPP1* (Figure S11). In an *in vitro* oxLDL-induced foam cell model using mouse BMDMs, oxLDL lowered *C1qb*/*C1qc* while raising *Mmp12*, *Pkm*, *Cd36*, plus *Prdx6*/*Hmox1* (Figure 6F). KEGG analysis showed oxLDL-induced genes in mineral absorption, glutathione, ferroptosis pathways, downregulated in antigen processing (Figure 6G). Western blotting analysis confirmed dose-dependent ALOX5, ALOX5AP, ACSL4 rises (Figure 6H), backing arachidonic metabolism and peroxidation^20^.

**Figure 6.**
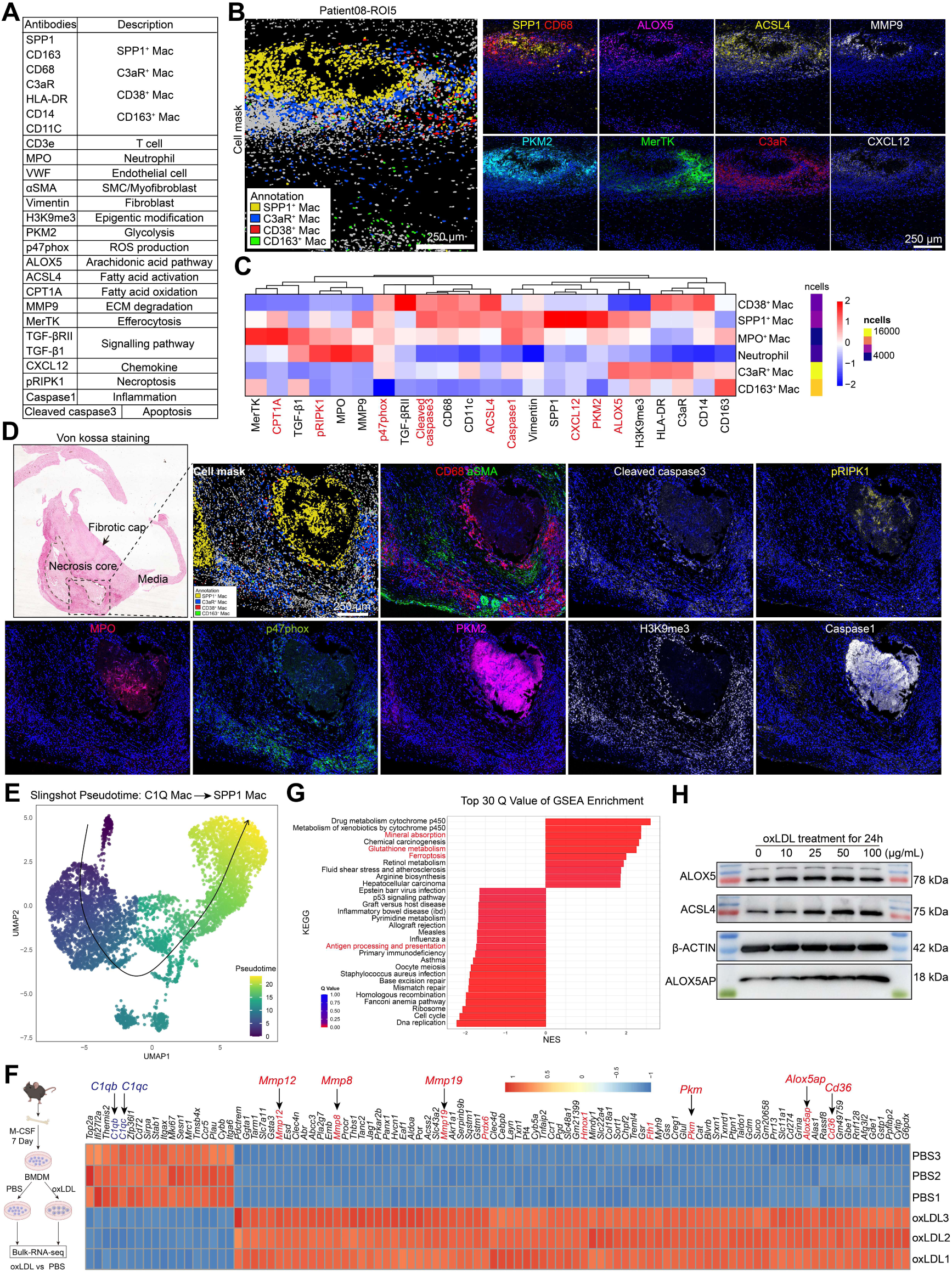
Functional profiling reveals distinct metabolic and death programs in macrophage subsets. **A,** List of antibodies included in IMC panel 2, targeting markers related to macrophage identity, efferocytosis, metabolism, ROS production, apoptosis, necroptosis, inflammation, epigenetic modification, chemokines (CXCL12), and fibroblast and smooth muscle cell markers (vimentin, αSMA). **B,** Representative IMC images from ROI5 showing spatial localization and marker expression in SPP1^+^, C3aR^+^, CD38^+^ and CD163^+^ macrophages. Scale bar, 250 µm. **C,** Heatmap showing expression levels of functional markers across macrophage subsets and other cell types, highlighting metabolic, oxidative, death, and epigenetic signatures. **D,** Von Kossa staining (far left) of advanced carotid plaque with adjacent IMC panels showing SPP1^+^ and C3aR^+^ macrophage distribution, and marker expression of CD68, αSMA, Cleaved caspase-3, pRIPK1, MPO, p47phox, PKM2, H3K9me3, and Caspase-1. Scale bar, 250 µm. **E,** Pseudotime analysis using Slingshot on public scRNA-seq data identified a continuous trajectory from C1Q macrophages toward SPP1 macrophages. Each dot represents a single macrophage, colored by pseudotime progression from early (blue) to late (yellow) stages. The black curve indicates the inferred developmental trajectory. **F,** Bulk RNA-seq analysis of oxLDL-treated mouse bone marrow-derived macrophages (BMDMs) showing downregulation of *C1qb* and *C1qc* and upregulation of foam cell-associated genes (e.g., *Mmp12*, *Mmp8*, *Mmp19*, *Pkm*, *Alox5ap*, *Cd36*). **G,** GSEA enrichment plot showing upregulated pathways and downregulated pathways in oxLDL-treated BMDMs compared to PBS control. **H,** Western blot validation showing dose-dependent upregulation of ALOX5, ALOX5AP, and ACSL4 proteins in BMDMs treated with oxLDL for 24 h.

Prior studies noted TREM2^hi^ macrophages shift to inflammatory PLIN2^hi^/TREM1^hi^ in human atherosclerosis^33^. Our scRNA-seq and immunohistochemistry showed C1Q/SPP1 macrophages express TREM2, HMOX-1, PLIN2 (Figure S2C and S9B-C); IMC had highest PLIN2 in SPP1 macrophage (MMP9^hi^) near cores, lower in C3aR^+^/CD38^+^ (Figure S9D). High HMOX1 areas had nitrotyrosine and macrophage density (Figure S9E), showing ROS/ONOOH in plaques with macrophages as oxidants. This outline reprogramming from C1Q to death-prone SPP1 (PLIN2^hi^/TREM1^hi^) macrophage in necrotic niches, though fate mapping needed for proof.

### C3aR^+^ macrophages create antigen-presenting niches in advanced plaques

High HLA-DR expression and antigen-presenting potential in C3aR^+^ macrophages suggest T cell interactions in plaques^39^. IMC showed them ringed by CD4^+^/CD8^+^ T cells, some FoxP3^+^ Tregs, and nearby mast cells (Figure 7A-B), indicating structured immune niches. Cellular neighborhood (CN) analysis found distinct microenvironments (Figure 7C-D). Key: CN1 (C3aR Mac_T) and CN11 (APC_T): Mostly C3aR^+^/CD38^+^ macrophages, HLA-DR^+^ SMCs, CD4^+^/CD8^+^ T cells. Macrophages/SMCs expressed high HLA-DR, as APCs engaging T cells. CN5 (Foamy_proliferating): SPP1^+,^ Ki67^+^, FN^+^ macrophages, for proliferation and remodeling, potentially contributing to necrotic core growth. CN9 (Endothelium): TGF-β1/pp65^+^ ECs, for endothelial-mesenchymal shift. CN10 (Media): Mainly SMCs, matching vascular media. Cell interaction analysis showed stronger C3aR^+^/CD38^+^ links to CD4^+^/CD8^+^ T cells than SPP1^+^ in advanced plaques (Figure 7E). Though SPP1^+^ abundant in necrotic cores, limited presentation may depend on C3aR^+^/CD38^+^ for ApoB epitopes to T cells^40,41^. Advanced plaques showed interactions between C3aR^+^ macrophages and fibroblasts; SPP1^+^ macrophages and Ki67^+^ or FN^+^ macrophages; CD4^+^ T cells and fibroblasts; as well as endothelial cells with CD4^+^ T cells and neutrophils. Ligand–receptor analysis focusing on all cells located in necrotic core regions identified CXCL12-CXCR4 signaling as a key axis mediating macrophage-T cell interactions, as well as interactions between mast cells and T cells, and between fibroblasts and endothelial cells with T cells (Figure 7F). This chemokine-mediated crosstalk may facilitate recruitment and retention of T cells within macrophage-rich niches, thereby promoting local immune activation^42,43^.

**Figure 7.**
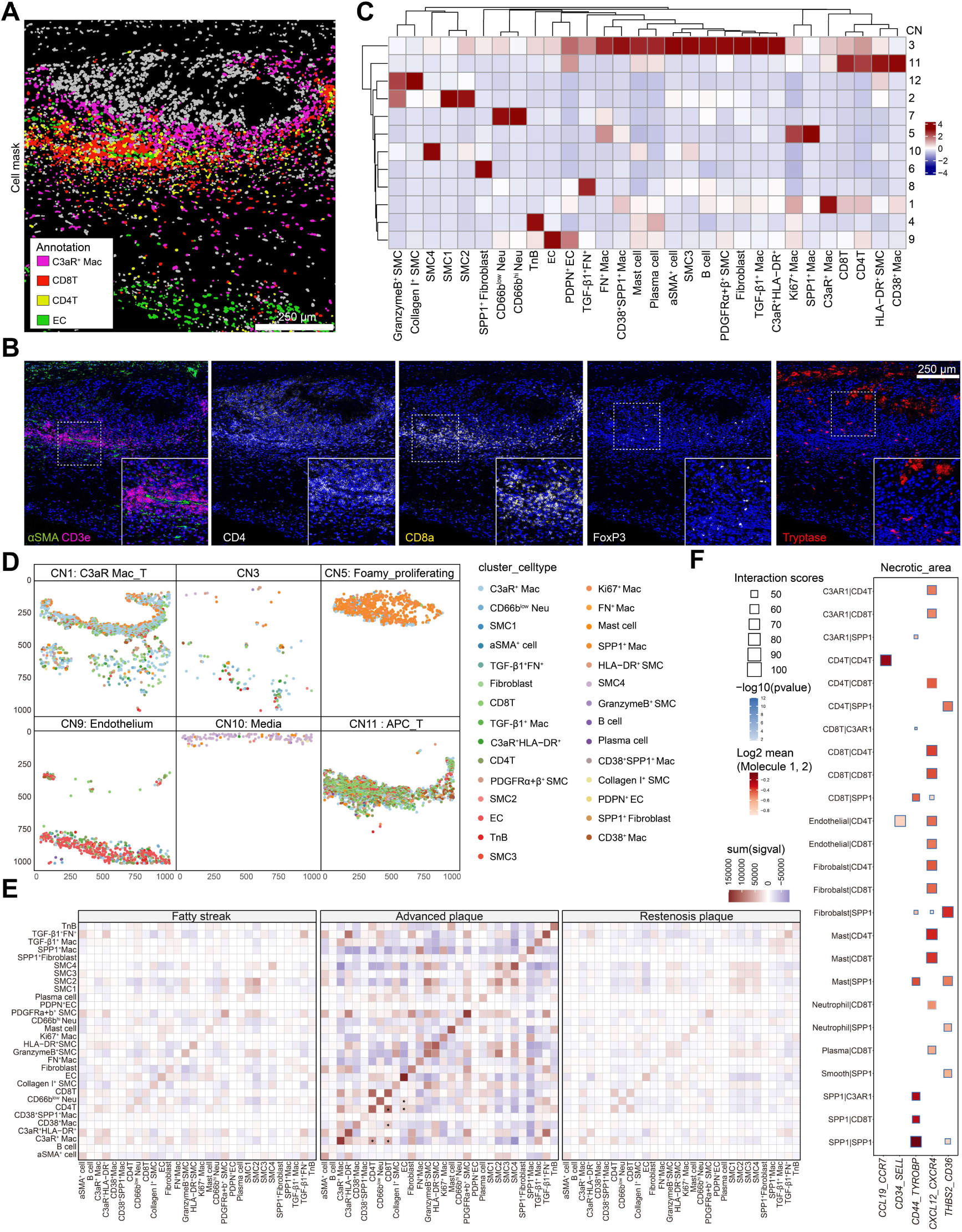
C3aR^+^ macrophages form antigen-presenting niches interacting with T cells in advanced plaques. **A,** Cell mask segmentation showing spatial localization of C3aR^+^ macrophages (rose purple), CD4^+^ T cells (orange), CD8^+^ T cells (red), and endothelial cells (green) within advanced plaque tissue. Scale bar, 250 µm. **B,** Representative IMC images showing marker expression for αSMA, CD3e, CD4, CD8α, FoxP3, and Mast cell tryptase. **C,** Dot plot showing relative frequency of major immune and stromal cell types across 10 defined cellular neighborhoods (CNs) within advanced plaques. **D,** Distribution of CNs in patient8-ROI5, highlighting key regions including CN1 (C3aR_Mac), CN5 (Foamy_proliferating), CN9 (Endothelium), CN10 (Media), and CN11 (APC_T) with cell type annotations. **E,** Heatmaps showing cell interaction scores among cell types within fatty streak, advanced, and restenotic plaques. In the plot above the red tiles indicate cell type pairs that were detected to significantly interact on a large number of images. Blue tiles show cell type pairs which tend to avoid each other on a large number of images. **F,** Ligand–receptor analysis focusing on necrotic core regions. Scale bar, 250 µm.

Together, these results reveal that C3aR^+^ macrophages establish specialized antigen-presenting niches, where enriched T cells in advanced plaques receive activation signals, supporting a model in which spatially organized myeloid–lymphoid interactions coordinate immune activation, inflammation, and tissue remodeling in human atherosclerosis.

## Discussion

Atherosclerosis advances through dynamic interactions among immune cells that influence plaque development and stability^44–46^. Current molecular profiling of cell types in plaques mainly depends on transcriptomic information^47,48^, even though RNA levels do not always match protein levels^49,50^. In this study, we combined high-resolution Visium HD spatial transcriptomics, imaging mass cytometry, and single-cell RNA sequencing from human fatty streak, advanced, and restenotic plaques to build a stage-specific immune atlas that tracks the disease’s progression. This approach uncovered specialized neutrophil and macrophage populations in distinct spatial locations, underscoring the role of tissue structure in determining immune roles in atherosclerotic plaques.

In particular, unlike a previous scRNA-seq-based atlas^33^ that detected 12 detailed myeloid groups such as C1Q^+^, HMOX1^+^, TREM2^hi^, and PLIN2^hi^/TREM1^hi^ macrophages, our imaging mass cytometry method pinpointed macrophage states fixed in space directly within human plaques. Due to the restricted number of metal channels and the lack of specificity of CD11c as a marker for dendritic cells in imaging mass cytometry, we avoided labeling dendritic cell subgroups. However, we found groups matching scRNA-seq clusters, like C1Q (C3aR^+^ macrophages), S100A8/9^+^ (neutrophils), and PLIN2^hi^/TREM1^hi^ (SPP1 macrophages). In line with scRNA-seq trajectory studies indicating that TREM2^hi^ macrophages evolve into an inflammatory PLIN2^hi^/TREM1^hi^ state, we noted the spatial overlap of PLIN2^hi^ and SPP1^+^ macrophages close to necrotic centers. On the other hand, areas with low PLIN2 were associated with antigen-presenting C3aR^+^ and pro-inflammatory CD38^+^ macrophages. These location-based patterns imply that C1Q (HMOX1^+^) macrophages could transform into the SPP1 macrophages, backed by in vitro reprogramming triggered by oxLDL. Crucially, identifying neutrophils was more robust with IMC than with scRNA-seq, illustrating the added value of spatial protein analysis compared to RNA methods.

In fatty streak lesions, neutrophils were dominant, producing oxidative (MPO, ALOX5, HMOX1) and proteolytic factors (including MMP9) that spark inflammation and tissue restructuring. Interestingly, these neutrophils aligned with fibronectin-abundant zones and spots of chlorotyrosine buildup, indicating that their oxidative and proteolytic actions help drive early extracellular matrix damage. By comparison, advanced plaques featured macrophage-led oxidative stress, as SPP1^+^ and C3aR^+^ groups showed strong levels of HMOX1 and p47phox, matching widespread nitrotyrosine presence. These observations point to a shift in oxidative stress origins from neutrophils in initial lesions to macrophages in later stages.

From a clinical view, the strong buildup of nitrotyrosine in unstable areas underscores its promise as an indicator of plaque weakness, connecting redox disruptions to the lesion’s structural breakdown. In advanced plaques, Ki67^+^ macrophages multiplied near necrotic centers^22^, aiding local population renewal and aligning with earlier reports that proliferation fuels macrophage buildup in mature plaques. Their proximity to SPP1^+^ macrophages implies that necrotic-core surroundings, rich in dead cell remnants, could trigger reparative macrophage growth. SPP1^+^ macrophages, concentrated near necrotic cores, also expressed glycolytic (PKM2), oxidative burst (p47phox, ALOX5, HMOX1), extracellular matrix restructuring (MMP9, Vimentin, Fibronectin), and markers for apoptosis or ferroptosis vulnerability^51,52^. PKM2, a key rate-limiting enzyme in glycolysis, rises in monocytes and macrophages from coronary atherosclerosis patients, where its levels strongly link to plaque instability^45^. Deletion of PKM2 from myeloid cells lowers macrophage death while boosting debris clearance, thus slowing atherosclerosis advancement. These patterns match our detection of high PKM2 in SPP1^+^ macrophages, pointing to a preference for glycolytic metabolism. Moreover, their elevated cleaved caspase-3 and reduced MerTK levels signal poor debris-handling ability and a tendency for cell death, which could enlarge necrotic cores and heighten plaque risk. Additionally, fibronectin buildup and its broken pieces, tied closely associated with SPP1^+^ macrophages, might serve as damage-associated molecular patterns (DAMPs) that intensify inflammatory pathways and worsen macrophage demise, thus tying matrix changes to further necrosis. Although foamy macrophages with high Trem2 and Spp1 expression have been previously reported^10,53^, our results combine metabolic, restructuring, and death-related traits into a fully mature foam cell-like condition, which likely drives necrotic core growth through matrix breakdown and controlled cell death, aiding plaque weakening.

Fibronectin (FN) accumulates at necrotic-core rims with SPP1⁺ macrophages, suggesting an ECM cue shaping their identity. During monocyte-to-macrophage differentiation, FN upregulated MMP9, TIMP1 and HK2, whereas FN applied to mature THP-1 macrophages had minimal effects, indicating stage-dependent imprinting. This matrix route complements oxLDL– driven lipid loading, with both converging on an SPP1-linked remodeling state. Interestingly, C3aR^+^ macrophages were rich in HLA-DR, CD74, and C1QC, creating antigen-presenting zones encircled by CD4^+^ T and CD8^+^ T cells. While macrophages are known as antigen presenters, spotting a C3aR^+^ macrophage-T cell zone is new in human plaques. Interaction predictions highlighted CXCL12–CXCR4 signaling as a link between them, implying chemokine-guided T cell retention and activation in these zones, which might boost nearby adaptive immune activity. Further, CD38^+^ macrophages had raised pp65 level, indicating NF-κB pathway engagement and an inflammatory profile. Though without clear spatial positioning, their presence in APC-T cell areas suggests they help spread general plaque inflammation together with antigen-presenting macrophages.

Our work builds on earlier single-cell and CyTOF studies in mouse and human plaques^8,11,20^, which defined macrophage diversity but overlooked their spatial arrangement ^10,54,55^. By merging near single-cell spatial transcriptomics with multi-protein imaging, we created a multi-omics overview of the human plaque myeloid setting, showing how groups are placed to manage matrix restructuring, lipid management, antigen display, and growth. Additionally, the buildup of H3K9me3 in SPP1^+^ and C3aR^+^ macrophages points to possible epigenetic control of their specialized roles, calling for deeper mechanistic exploration.

Drawbacks involve using carotid or femoral fixed samples, which might vary from coronary ones, and the mostly observational style of our work. Upcoming studies should add functional disruption methods, spatial epigenetics, and live cell tracking to clarify how surrounding and epigenetic signals shape macrophage roles, foam cell demise, and antigen display. To strengthen causality, receptor-level tests (α5β1 blockade, cRGD competition, FAK/PI3K inhibition) and matrix-mechanics manipulations (FN fiber assembly, substrate stiffness, EDA⁺ vs EDA⁻ isoforms) in primary human macrophages will be informative and clarify interplay with lipid-uptake pathways. Unraveling these routes could uncover treatments to adjust targeted myeloid zones, curb inflammation, and secure unstable plaques.

Overall, this spatial myeloid atlas shows that the layout of neutrophil and macrophage activities strongly guides their roles in atherosclerosis. Through stage-focused myeloid zones, our results deepen insights into plaque immune pathology and offer a basis for therapies aimed at the microenvironment to secure plaques and lower heart disease threats.

## Acknowledgements

We thank the Bodenmiller Laboratory for generously providing their imaging mass cytometry data toolkit and IMC-DATA analysis workflow, and the Peng Lu Laboratory for developing the IMC_Denoise tool. Finally, we acknowledge Cosmos Wisdom Biotech Co., Ltd., Dr. Zhentao Song and Dr. Taotao Zhu for providing 10x Genomics Visium HD spatial transcriptomics sequencing and data analysis support.

## Author Contributions

P.H. conducted the majority of the IMC experiments, processed computational data, and prepared the figures. B.H. contributed clinical expertise, patient samples, and associated medical data. P.H., Z.L., S.W., and Z.W. performed antibody conjugation and validation; P.H. and Z.W. carried out histopathological staining. Y.S., K.L., Q.L., H.C., and J.F. provided constructive feedback during manuscript preparation. K.L., P.L., G.M., and C.S. offered project guidance and manuscript revisions. Y.S. oversaw study conception and design, manuscript writing, administrative support, and financial support.

## References

1. Libby P. The changing landscape of atherosclerosis. Nature. 2021;592:524–533. doi: 10.1038/s41586-021-03392-8

2. Herrington W, Lacey B, Sherliker P, Armitage J, Lewington S. Epidemiology of Atherosclerosis and the Potential to Reduce the Global Burden of Atherothrombotic Disease. Circ Res. 2016;118:535–546. doi: 10.1161/CIRCRESAHA.115.307611

3. Martin SS, Aday AW, Almarzooq ZI, Anderson CAM, Arora P, Avery CL, Baker-Smith CM, Barone Gibbs B, Beaton AZ, Boehme AK, et al. 2024 Heart Disease and Stroke Statistics: A Report of US and Global Data From the American Heart Association. Circulation. 2024;149:e347–e913. doi: 10.1161/CIR.0000000000001209

4. Vaduganathan M, Mensah GA, Turco JV, Fuster V, Roth GA. The Global Burden of Cardiovascular Diseases and Risk: A Compass for Future Health. J Am Coll Cardiol. 2022;80:2361–2371. doi: 10.1016/j.jacc.2022.11.005

5. Ridker PM, MacFadyen JG, Everett BM, Libby P, Thuren T, Glynn RJ, Group CT. Relationship of C-reactive protein reduction to cardiovascular event reduction following treatment with canakinumab: a secondary analysis from the CANTOS randomised controlled trial. Lancet. 2018;391:319–328. doi: 10.1016/S0140-6736(17)32814-3

6. Ridker PM, Devalaraja M, Baeres FMM, Engelmann MDM, Hovingh GK, Ivkovic M, Lo L, Kling D, Pergola P, Raj D, et al. IL-6 inhibition with ziltivekimab in patients at high atherosclerotic risk (RESCUE): a double-blind, randomised, placebo-controlled, phase 2 trial. Lancet. 2021;397:2060–2069. doi: 10.1016/S0140-6736(21)00520-1

7. Fernandez DM, Giannarelli C. Immune cell profiling in atherosclerosis: role in research and precision medicine. Nat Rev Cardiol. 2022;19:43–58. doi: 10.1038/s41569-021-00589-2

8. Fernandez DM, Rahman AH, Fernandez NF, Chudnovskiy A, Amir ED, Amadori L, Khan NS, Wong CK, Shamailova R, Hill CA, et al. Single-cell immune landscape of human atherosclerotic plaques. Nat Med. 2019;25:1576–1588. doi: 10.1038/s41591-019-0590-4

9. Traeuble K, Munz M, Pauli J, Sachs N, Vafadarnejad E, Carrillo-Roa T, Maegdefessel L, Kastner P, Heinig M. Integrated single-cell atlas of human atherosclerotic plaques. Nat Commun. 2025;16:8255. doi: 10.1038/s41467-025-63202-x

10. Cochain C, Vafadarnejad E, Arampatzi P, Pelisek J, Winkels H, Ley K, Wolf D, Saliba AE, Zernecke A. Single-Cell RNA-Seq Reveals the Transcriptional Landscape and Heterogeneity of Aortic Macrophages in Murine Atherosclerosis. Circ Res. 2018;122:1661–1674. doi: 10.1161/CIRCRESAHA.117.312509

11. Winkels H, Ehinger E, Vassallo M, Buscher K, Dinh HQ, Kobiyama K, Hamers AAJ, Cochain C, Vafadarnejad E, Saliba AE, et al. Atlas of the Immune Cell Repertoire in Mouse Atherosclerosis Defined by Single-Cell RNA-Sequencing and Mass Cytometry. Circ Res. 2018;122:1675–1688. doi: 10.1161/CIRCRESAHA.117.312513

12. Hou P, Fang J, Liu Z, Shi Y, Agostini M, Bernassola F, Bove P, Candi E, Rovella V, Sica G, et al. Macrophage polarization and metabolism in atherosclerosis. Cell Death Dis. 2023;14:691. doi: 10.1038/s41419-023-06206-z

13. Silvestre-Roig C, Braster Q, Ortega-Gomez A, Soehnlein O. Neutrophils as regulators of cardiovascular inflammation. Nat Rev Cardiol. 2020;17:327–340. doi: 10.1038/s41569-019-0326-7

14. Soehnlein O. Multiple roles for neutrophils in atherosclerosis. Circ Res. 2012;110:875–888. doi: 10.1161/CIRCRESAHA.111.257535

15. Armingol E, Officer A, Harismendy O, Lewis NE. Deciphering cell-cell interactions and communication from gene expression. Nat Rev Genet. 2021;22:71–88. doi: 10.1038/s41576-020-00292-x

16. Bleckwehl T, Babler A, Tebens M, Maryam S, Nyberg M, Bosteen M, Halder M, Shaw I, Fleig S, Pyke C, et al. Encompassing view of spatial and single-cell RNA sequencing renews the role of the microvasculature in human atherosclerosis. Nat Cardiovasc Res. 2025;4:26–44. doi: 10.1038/s44161-024-00582-1

17. Chang Q, Ornatsky OI, Siddiqui I, Loboda A, Baranov VI, Hedley DW. Imaging Mass Cytometry. Cytometry A. 2017;91:160–169. doi: 10.1002/cyto.a.23053

18. Hartmann FJ, Bendall SC. Immune monitoring using mass cytometry and related high-dimensional imaging approaches. Nat Rev Rheumatol. 2020;16:87–99. doi: 10.1038/s41584-019-0338-z

19. Bressan D, Battistoni G, Hannon GJ. The dawn of spatial omics. Science. 2023;381:eabq4964. doi: 10.1126/science.abq4964

20. Zernecke A, Erhard F, Weinberger T, Schulz C, Ley K, Saliba AE, Cochain C. Integrated single-cell analysis-based classification of vascular mononuclear phagocytes in mouse and human atherosclerosis. Cardiovasc Res. 2023;119:1676–1689. doi: 10.1093/cvr/cvac161

21. Sharifi MA, Wierer M, Dang TA, Milic J, Moggio A, Sachs N, von Scheidt M, Hinterdobler J, Muller P, Werner J, et al. ADAMTS-7 Modulates Atherosclerotic Plaque Formation by Degradation of TIMP-1. Circ Res. 2023;133:674–686. doi: 10.1161/CIRCRESAHA.123.322737

22. Robbins CS, Hilgendorf I, Weber GF, Theurl I, Iwamoto Y, Figueiredo JL, Gorbatov R, Sukhova GK, Gerhardt LM, Smyth D, et al. Local proliferation dominates lesional macrophage accumulation in atherosclerosis. Nat Med. 2013;19:1166–1172. doi: 10.1038/nm.3258

23. Terao R, Lee TJ, Colasanti J, Pfeifer CW, Lin JB, Santeford A, Hase K, Yamaguchi S, Du D, Sohn BS, et al. LXR/CD38 activation drives cholesterol-induced macrophage senescence and neurodegeneration via NAD(+) depletion. Cell Rep. 2024;43:114102. doi: 10.1016/j.celrep.2024.114102

24. Covarrubias AJ, Kale A, Perrone R, Lopez-Dominguez JA, Pisco AO, Kasler HG, Schmidt MS, Heckenbach I, Kwok R, Wiley CD, et al. Senescent cells promote tissue NAD(+) decline during ageing via the activation of CD38(+) macrophages. Nat Metab. 2020;2:1265–1283. doi: 10.1038/s42255-020-00305-3

25. Ma Y, Zhang Y, Liu X, Yang X, Guo H, Ding X, Ye C, Guo C. Deletion of CD38 mitigates the severity of NEC in experimental settings by modulating macrophage-mediated inflammation. Redox Biol. 2024;77:103336. doi: 10.1016/j.redox.2024.103336

26. Chen Y, Yang M, Huang W, Chen W, Zhao Y, Schulte ML, Volberding P, Gerbec Z, Zimmermann MT, Zeighami A, et al. Mitochondrial Metabolic Reprogramming by CD36 Signaling Drives Macrophage Inflammatory Responses. Circ Res. 2019;125:1087–1102. doi: 10.1161/CIRCRESAHA.119.315833

27. Park YM. CD36, a scavenger receptor implicated in atherosclerosis. Exp Mol Med. 2014;46:e99. doi: 10.1038/emm.2014.38

28. Slysz J, Sinha A, DeBerge M, Singh S, Avgousti H, Lee I, Glinton K, Nagasaka R, Dalal P, Alexandria S, et al. Single-cell profiling reveals inflammatory polarization of human carotid versus femoral plaque leukocytes. JCI Insight. 2023;8. doi: 10.1172/jci.insight.171359

29. Souilhol C, Harmsen MC, Evans PC, Krenning G. Endothelial-mesenchymal transition in atherosclerosis. Cardiovasc Res. 2018;114:565–577. doi: 10.1093/cvr/cvx253

30. Andueza A, Kumar S, Kim J, Kang DW, Mumme HL, Perez JI, Villa-Roel N, Jo H. Endothelial Reprogramming by Disturbed Flow Revealed by Single-Cell RNA and Chromatin Accessibility Study. Cell Rep. 2020;33:108491. doi: 10.1016/j.celrep.2020.108491

31. Yang-Jensen KC, Jorgensen SM, Chuang CY, Davies MJ. Modification of extracellular matrix proteins by oxidants and electrophiles. Biochem Soc Trans. 2024;52:1199–1217. doi: 10.1042/BST20230860

32. Nybo T, Cai H, Chuang CY, Gamon LF, Rogowska-Wrzesinska A, Davies MJ. Chlorination and oxidation of human plasma fibronectin by myeloperoxidase-derived oxidants, and its consequences for smooth muscle cell function. Redox Biol. 2018;19:388–400. doi: 10.1016/j.redox.2018.09.005

33. Dib L, Koneva LA, Edsfeldt A, Zurke YX, Sun J, Nitulescu M, Attar M, Lutgens E, Schmidt S, Lindholm MW, et al. Lipid-associated macrophages transition to an inflammatory state in human atherosclerosis increasing the risk of cerebrovascular complications. Nat Cardiovasc Res. 2023;2:656–672. doi: 10.1038/s44161-023-00295-x

34. de Gaetano G, Donati MB, Cerletti C. Prevention of thrombosis and vascular inflammation: benefits and limitations of selective or combined COX-1, COX-2 and 5-LOX inhibitors. Trends Pharmacol Sci. 2003;24:245–252. doi: 10.1016/S0165-6147(03)00077-4

35. Gilbert NC, Gerstmeier J, Schexnaydre EE, Borner F, Garscha U, Neau DB, Werz O, Newcomer ME. Structural and mechanistic insights into 5-lipoxygenase inhibition by natural products. Nat Chem Biol. 2020;16:783–790. doi: 10.1038/s41589-020-0544-7

36. Chen S, Yang J, Wei Y, Wei X. Epigenetic regulation of macrophages: from homeostasis maintenance to host defense. Cell Mol Immunol. 2020;17:36–49. doi: 10.1038/s41423-019-0315-0

37. Groh L, Keating ST, Joosten LAB, Netea MG, Riksen NP. Monocyte and macrophage immunometabolism in atherosclerosis. Semin Immunopathol. 2018;40:203–214. doi: 10.1007/s00281-017-0656-7

38. Kuznetsova T, Prange KHM, Glass CK, de Winther MPJ. Transcriptional and epigenetic regulation of macrophages in atherosclerosis. Nat Rev Cardiol. 2020;17:216–228. doi: 10.1038/s41569-019-0265-3

39. Mohanta SK, Heron C, Klaus-Bergmann A, Horstmann H, Brakenhielm E, Giannarelli C, Habenicht AJR, Gerhardt H, Weber C. Metabolic and Immune Crosstalk in Cardiovascular Disease. Circ Res. 2025;136:1433–1453. doi: 10.1161/CIRCRESAHA.125.325496

40. Wolf D, Ley K. Immunity and Inflammation in Atherosclerosis. Circ Res. 2019;124:315–327. doi: 10.1161/CIRCRESAHA.118.313591

41. Saigusa R, Winkels H, Ley K. T cell subsets and functions in atherosclerosis. Nat Rev Cardiol. 2020;17:387–401. doi: 10.1038/s41569-020-0352-5

42. Doring Y, Noels H, van der Vorst EPC, Neideck C, Egea V, Drechsler M, Mandl M, Pawig L, Jansen Y, Schroder K, et al. Vascular CXCR4 Limits Atherosclerosis by Maintaining Arterial Integrity: Evidence From Mouse and Human Studies. Circulation. 2017;136:388–403. doi: 10.1161/CIRCULATIONAHA.117.027646

43. Doring Y, van der Vorst EPC, Weber C. Targeting immune cell recruitment in atherosclerosis. Nat Rev Cardiol. 2024;21:824–840. doi: 10.1038/s41569-024-01023-z

44. Roy P, Orecchioni M, Ley K. How the immune system shapes atherosclerosis: roles of innate and adaptive immunity. Nat Rev Immunol. 2022;22:251–265. doi: 10.1038/s41577-021-00584-1

45. Palm KCA, Yin X, Baig F, Theofilatos K, van der Laan SW, de Borst GJ, de Kleijn DPV, Wojta J, Stojkovic S, Mayr M, et al. Proteomic profiling reveals a higher presence of glycolytic enzymes in human atherosclerotic lesions with unfavourable histological characteristics. Cardiovasc Res. 2025. doi: 10.1093/cvr/cvaf077

46. Gianopoulos I, Daskalopoulou SS. Macrophage profiling in atherosclerosis: understanding the unstable plaque. Basic Res Cardiol. 2024;119:35–56. doi: 10.1007/s00395-023-01023-z

47. Vallejo J, Cochain C, Zernecke A, Ley K. Heterogeneity of immune cells in human atherosclerosis revealed by scRNA-Seq. Cardiovasc Res. 2021;117:2537–2543. doi: 10.1093/cvr/cvab260

48. Zernecke A, Winkels H, Cochain C, Williams JW, Wolf D, Soehnlein O, Robbins CS, Monaco C, Park I, McNamara CA, et al. Meta-Analysis of Leukocyte Diversity in Atherosclerotic Mouse Aortas. Circ Res. 2020;127:402–426. doi: 10.1161/CIRCRESAHA.120.316903

49. Upadhya SR, Ryan CJ. Experimental reproducibility limits the correlation between mRNA and protein abundances in tumor proteomic profiles. Cell Rep Methods. 2022;2:100288. doi: 10.1016/j.crmeth.2022.100288

50. Liu Y, Beyer A, Aebersold R. On the Dependency of Cellular Protein Levels on mRNA Abundance. Cell. 2016;165:535–550. doi: 10.1016/j.cell.2016.03.014

51. Gerlach BD, Ampomah PB, Yurdagul A, Jr., Liu C, Lauring MC, Wang X, Kasikara C, Kong N, Shi J, Tao W, Tabas I. Efferocytosis induces macrophage proliferation to help resolve tissue injury. Cell Metab. 2021;33:2445–2463 e2448. doi: 10.1016/j.cmet.2021.10.015

52. Ngai D, Schilperoort M, Tabas I. Efferocytosis-induced lactate enables the proliferation of pro-resolving macrophages to mediate tissue repair. Nat Metab. 2023;5:2206–2219. doi: 10.1038/s42255-023-00921-9

53. Mosquera JV, Auguste G, Wong D, Turner AW, Hodonsky CJ, Alvarez-Yela AC, Song Y, Cheng Q, Lino Cardenas CL, Theofilatos K, et al. Integrative single-cell meta-analysis reveals disease-relevant vascular cell states and markers in human atherosclerosis. Cell Rep. 2023;42:113380. doi: 10.1016/j.celrep.2023.113380

54. Cole JE, Park I, Ahern DJ, Kassiteridi C, Danso Abeam D, Goddard ME, Green P, Maffia P, Monaco C. Immune cell census in murine atherosclerosis: cytometry by time of flight illuminates vascular myeloid cell diversity. Cardiovasc Res. 2018;114:1360–1371. doi: 10.1093/cvr/cvy109

55. Depuydt MAC, Prange KHM, Slenders L, Ord T, Elbersen D, Boltjes A, de Jager SCA, Asselbergs FW, de Borst GJ, Aavik E, et al. Microanatomy of the Human Atherosclerotic Plaque by Single-Cell Transcriptomics. Circ Res. 2020;127:1437–1455. doi: 10.1161/CIRCRESAHA.120.316770

